# *In silico* characterization of five novel disease-resistance proteins in *Oryza sativa sp. japonica* against bacterial leaf blight and rice blast diseases

**DOI:** 10.1101/2023.06.05.543715

**Authors:** Vedikaa Dhiman, Soham Biswas, Rajveer Singh Shekhawat, Ayan Sadhukhan, Pankaj Yadav

**Author notes:** **Correspondence:** Pankaj Yadav, Assistant Professor, Department of Bioscience and Bioengineering, Indian Institute of Technology, Jodhpur, 342030 Rajasthan, India.

## Abstract

*Oryza sativa* sp. *japonica* is the most widely cultivated variety of rice. It has evolved several defense mechanisms, including PAMP-triggered immunity (PTI) and effector-triggered immunity (ETI), which provide resistance against different pathogens to overcome biotic stresses. Several disease-resistance genes and proteins, such as R genes and PRR proteins, have been reported in the scientific literature which shows resistance against *Xanthomonas oryzae* pv. *oryzae* (*Xoo*), a causative agent for bacterial leaf blight disease (BB), and *Magnaporthe oryzae* (*M. oryzae*), causing rice blast disease (RB). Although some of these resistance proteins have been studied, the functional characterization of resistance proteins in rice is not exhaustive. In the current study, we identified five novel resistance proteins against BB and RB diseases through gene network analysis. Structure and function prediction, disease-resistance domain identification, protein-protein interaction (PPI), and pathway analysis revealed that the five new proteins played a role in the disease resistance against BB and RB. *In silico* modeling, refinement, and model quality assessment were performed to predict the best structures of these five proteins, and submitted to ModelArchive for future use. The functional annotation of the proteins revealed their involvement in the bacterial disease resistance of rice. We predicted that the new resistance proteins could be localized to the nucleus and plasma membrane. This study provides insight into developing disease-resistant rice varieties by predicting novel candidate resistance proteins, which will pave the way for their future characterization and assist rice breeders in improving crop yield and addressing future food security.

## INTRODUCTION

Rice (*Oryza sativa* sp. *japonica*) is a monocotyledonous angiosperm whose genome comprises 12 chromosomes. It is consumed by over 90% of the world’s population but produced by only a few countries. The top rice-producing countries, China, India, and Indonesia, produce 30.85%, 20.12%, and 8.21% of total global rice production, respectively (S. Kumar et al., 2017). By 2050, the demand for food and fiber will rise by up to 70% globally (Singh & Trivedi, 2017). This increasing demand for agricultural production must be achieved in existing arable land, under harsher climatic conditions, with deteriorating soil and water quality.

Several pathogenic bacteria and fungi attack the *Oryza sativa* sp. *japonica* and massively reduce its production. In recent years, the use of pesticides has decreased bacterial and fungal disease incidence but, at the same time, adversely impacted human health and the entire ecosystem. Alternatively, using disease-resistant varieties of *Oryza sativa* sp. *japonica* is considered an effective and sustainable disease control approach. Therefore, extensive studies have been conducted in recent years on pathogen identification in rice and the signaling mechanism responsible for recognizing innate immunity (Gowda et al., 2015).

The biotic stressors, such as bacteria and fungi, are the primary source of infection in different parts of the rice plants. *Xoo* is a causative agent in BB disease. It is a major devastating disease in the *Oryza sativa* sp. *japonica*, affecting millions of hectares of rice annually, with an essential crop loss of as high as 75% (He et al., 2022). Recently, in vitro studies showed that plant growth-promoting rhizobacteria like *Bacillus pumilus* SE34 and *Bacillus subtilis* GBO3 induce systematic resistance against *Xoo* and improved nutrient uptake and yield (Chithrashree et al., 2011). Though the use of these rhizobacteria might be beneficial, the genetic selection of resistant rice cultivars is the most effective control method for BB disease (Bakade et al., 2021). Further, *M. oryzae* is another causative agent of a major fungal disease known as RB. This disease causes a severe threat to rice yield across many rice-producing countries all over the world. It has been reported that RB causes 10–30% of the world’s food loss each year (Qi et al., 2023). RB has also hampered cereal crop production worldwide due to its high genetic variability, significantly affecting rice breeders and pathologists (Li et al., 2017). Recent studies have identified resistance genes like *Pigm, Ptr, Pi65 (t)*, and *bsr-d1* to show strong resistance against RB disease without reducing the quality or yield of rice (Zhai et al., 2019). Moreover, several broad-spectrum pathogen-resistant varieties of *Oryza sativa* sp. *japonica* are now available. For instance, in 2017, some researchers found that the GY129 variety was resistant to all types of *M. oryzae* isolates and was predicted to carry a novel blast *R* gene in northern China. In addition, a gene *Pi65 (t)* in the GY129 variety provides resistance against RB disease in the *japonica* rice cultivar (Wang et al., 2016).

Plants have a variety of defense mechanisms against biotic stresses, including two types of the immune system. The first type consists of pattern recognition receptors (PRRs) that identify pathogen-associated molecular patterns (PAMPs) leading to PTI. The second type comprises highly polymorphic resistance proteins (i.e., R proteins) against effectors on the pathogen’s external surface that initiate ETI. Several experimental studies have been performed on disease-resistance genes and proteins in *Oryza sativa* sp. *japonica*. For instance, disease-resistant rice varieties were developed using molecular marker-assisted selection by transferring broad-spectrum rice blast resistance genes into susceptible genotypes (Mi et al., 2018). In another study, a Quantitative Train Loci (QTL), qBBR11-1, was discovered in a cross of Teqing and Lemont cultivars, showing two years of resistance to BB infection caused by three types of *Xoo*, i.e., C2, C4, and C510 (Kumar et al., 2018).

Despite the success of single gene-based studies, the role of other disease-resistance genes and their pathways remained unexplored. In this context, a comprehensive analysis of disease-resistance genes at the genome-wide level is lacking for *Oryza sativa* sp. *japonica*. In the present study, we identified novel resistance proteins and annotated the uncharacterized disease-resistance proteins against BB and RB diseases using state-of-the-art bioinformatics tools. We identified, characterized, and validated disease-resistance proteins for BB and RB through a genome-wide study. We annotated these proteins according to their functions, domains, protein-protein interactions, and involvement in relevant biological pathways.

## MATERIALS & METHODS

### Data collection

The nucleotide sequences of the 182,437 genes from 12 chromosomes of the *Oryza sativa* sp. *japonica* were retrieved from the Ensembl plants database using the Biomart tool (https://plants.ensembl.org/index.html). Of these, 177,317 genes were known, and 5,120 genes were labeled as hypothetically conserved. The SNP-Seek database (https://snp-seek.irri.org/) retrieved previously known disease-resistance genes, including 56 and 85 disease-resistance genes for BB and RB diseases, respectively. **Table S1** provides an overview of the different tools and databases used in this study.

### Gene network construction and functional analysis

The gene network was constructed for hypothetically conserved genes and known disease-resistance genes using the Expasy STRING (Search Tool for the Retrieval of Interacting Genes/Proteins) tool (Snel et al., 2000). The STRING is the knowledgebase and software tool for known and predicted protein-protein interactions. Moreover, it includes direct (physical) and indirect (functional) associations derived from various sources, such as genomic context, high-throughput experiments, (conserved) co-expression, and the literature. The Cytoscape (version 3.9.1) tool was used for gene network analysis (Shannon et al., 2003). At first, the gene networks were separately exported for BB and RB from the Expasy STRING tool into the Cytoscape tool. Next, the genes (or nodes) in the network were colored differentially based on their function. The gene function information can be found in the ‘function’ column inside the node table available in the Cytoscape. Then, the first neighbors of each disease-resistance gene were selected, and the protein identifiers of these first neighbors were extracted from the node table. Only those proteins for which structural and functional information was unavailable in the UniProtKnowledgebase (UniProtKB) database were considered for further functional characterization.

### Physicochemical analyses of unknown disease-resistance proteins

The amino acid sequences of the unknown proteins identified above were retrieved from the UniProtKB database (Bateman et al., 2021). Furthermore, the retrieved amino acid sequences from the UniProtKB database were then subjected to validation for their various stereochemical and physicochemical properties. The physical and chemical properties, such as molecular weight, amino acid composition, theoretical isoelectric point, instability index, extinction coefficient, etc., of the identified unknown disease-resistance proteins, were determined using the ExPASy ProtParam tool (Gasteiger et al., 2003).

### Structure prediction and stereochemical property analysis of unknown disease-resistance proteins

The secondary structures of the unknown proteins were predicted using their amino acid sequences with PSI-BLAST Based Secondary Structure Prediction (PSIPRED) tool (McGuffin et al., 2000). This tool employs machine learning techniques to predict secondary structures for the input amino acid sequence. In addition, the 3D structures of these unknown proteins were predicted using SWISS-MODEL, a fully automated protein homology modeling server (Waterhouse et al., 2018). In this server, the template sequences for each protein were selected based on their Global Model Quality Estimate (GMQE) score and sequence identity. The GMQE score ranges from 0 to 1, with higher scores indicating more reliability of the template with the target sequence. In addition to the quality score, the sequence identity of the templates should be between 30% to 100%. The 3D structures predicted using SWISS-MODEL were further refined with the GalaxyRefine2 server (Lee et al., 2019) and ModRefiner (Xu & Zhang, 2011) and later visualized using PyMol (Schrödinger & DeLano, 2020). Furthermore, the neural network-based predictor CYSPRED (Fariselli et al., 1999) was used to determine the disulfide bonds between the cysteine residues of these disease-resistance proteins. It provides information regarding the number of cysteine residues, their bonding or non-bonding state, and their reliability score.

The refined protein structures obtained from the above analysis were then subjected to validation for their various stereochemical properties. The stereochemical properties of the proteins were verified using the Ramachandran plot server (Sheik et al., 2002) and the PROCHECK tool (Laskowski et al., 1993). The Ramachandran plot server was utilized for establishing the overall quality of the proteins, whereas PROCHECK studies the overall model geometry with the residue-by-residue geometry and provides the stereochemical quality of predicted models. Further, the quality of the protein models was evaluated using Verify3D (Eisenberg et al., 1997) and ERRAT (Colovos & Yeates, 1993). The Verify3D tool determines the compatibility of the atomic models with their own amino acid sequences. According to this tool, the percentage of compatibility should be greater than 80%. The ERRAT tool is a program for validating crystallography-determined protein structures in which a resolution of 95% or more is acceptable. Finally, all the predicted and refined structures of the unknown disease-resistance proteins identified in our study are submitted for public access at ModelArchive. The ModelArchive provides a unique stable accession code for each deposited model.

### Subcellular localization analysis

CELLO2GO web-based server was used to determine the subcellular localization of unknown disease-resistance proteins (Yu et al., 2014). In addition, the topology of these proteins was predicted using the HMMTOP server (Tusnády & Simon, 2001). HMMTOP transmembrane topology prediction server predicts both the localization of helical transmembrane segments and the topology of transmembrane proteins.

### Multiple sequence alignment and phylogenetic analysis

The protein family and domain information for the identified unknown disease-resistance proteins was retrieved from the InterProScan (Blum et al., 2021) and Conserved Domain Database (Lu et al., 2020), respectively. Furthermore, the disease-resistant domains were predicted in the identified proteins using the Leucine Rich Repeats (LRR) predictor (Martin et al., 2020) and Plant Resistance Genes Database (PRGdb) (Calle García et al., 2022). LRR predictor is a web server based on an ensemble of estimators designed to better identify LRR motifs. In the PRGdb, the DRAGO3 tool was used to predict disease-resistant domains and their positions. Additionally, we used ScanProsite (de Castro et al., 2006) and Motif Scan tools (Pagni et al., 2007) to find the consensus sequences and motifs for unknown disease-resistance proteins, respectively. The motifs identified using the Motif Scan tool were used further as input to ScanProsite for identifying the families of respective disease-resistance proteins. The amino acid sequences of the top five proteins from each identified family were retrieved for performing a BLAST search. The BLASTP tool was used to perform BLAST search taking each unknown disease-resistance protein and the identified top proteins from each family as input query against various databases such as UniProtKB, SwissProt, and TrEMBL. We selected the top 12 to 15 hits from the BLAST search with an alignment score >100 as a threshold, e-value close to 0, and similarity percentage greater than 60% for performing the local multiple sequence alignments (MSA). The *ggmsa* package (Zhou et al., 2022) in R software (version 4.2.2) was used to perform the MSA.

Furthermore, the phylogenetic analysis of the MSA results was performed using the neighbor-joining method available in the *phylogram* package (Wilkinson & Davy, 2018) in R software. The phylogenetic tree was constructed through bootstrapping for 100 cycles to get more statistically confident node distances. The phylogenetic trees were plotted using the *ggplot* package (Wickham, 2014) in R software.

### Functional enrichment analysis of disease-resistance proteins

All disease-resistance proteins were functionally annotated using PANNZER (Protein Annotation with Z-score) web server (Törönen & Holm, 2022). It provides information about protein functions in prokaryotic and eukaryotic organisms and gene ontology (GO) annotations. Furthermore, the pathway analysis was performed for the identified disease-resistance proteins to identify the presence or absence of a disease-resistance protein against *Xoo* and *M. oryzae*. For this purpose, the Assign KO tool from the KEGG Database (Kanehisa & Subramaniam, 2002) was used in the interactive mode. This tool is an interface to the BlastKOALA server, which assigns K numbers to the sequences provided by the user as input and performs a BLAST against a nonredundant set of KEGG genes.

### Identification of functional interactors in disease-resistance proteins

The proteins and their functional interactors play an important role in maintaining cellular processes. These interactors must be studied in order to gain a better understanding of various biological phenomena. Thus, to study the interaction of disease-resistance proteins with other similar proteins, the STRING database (Szklarczyk et al., 2019) was used. It is a biological database and web resource which incorporates all known and predicted physical and functional interactions between proteins.

## RESULTS

### Gene network analysis for hypothetically conserved and known disease-resistance genes

Gene networks were created by the Expasy STRING tool between hypothetically conserved and disease-resistance genes for BB and RB diseases (**Figure 1**). For BB disease, we have found 27 genes to be involved in disease resistance as well as in defense-related mechanisms. Similarly, for RB disease, we identified 26 genes engaged in disease resistance and defense-related mechanisms. The protein identifiers of the first neighbors of the disease-resistance genes were selected from the node table in Cytoscape. There were 125 and 129 as the first neighbors identified for BB and RB diseases, respectively. These 254 protein identifiers were further analyzed in the UniProtKB database. Of these, the UniProtKB database marked 5 proteins as unknown because these proteins do not have any structural and functional information available. For the purpose of this study, we named these 5 proteins according to their involvement in the disease. For instance, the 2 proteins involved in BB disease are Protein BB.1 and Protein BB.2. Similarly, 3 proteins involved in RB were named Protein RB.1, Protein RB.2, and Protein RB.3.

**Fig. 1.**
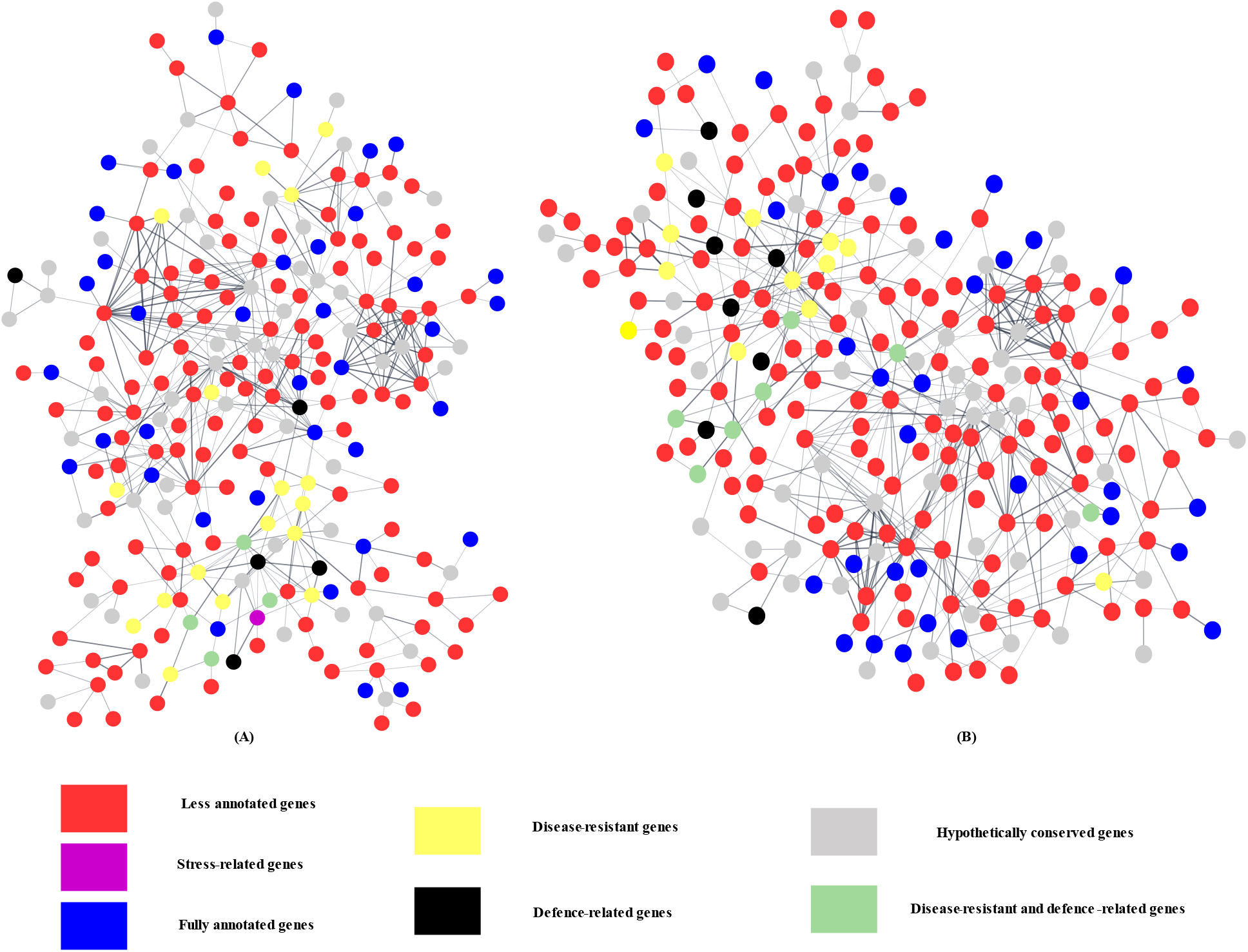
Cytoscape network of hypothetically conserved and known disease-resistance genes. **Panel (A):** A network between hypothetically conserved and bacterial blight disease-resistance genes. **Panel (B)**: A network between hypothetically conserved and rice blast disease-resistance genes. Here, yellow, black, green, purple, red, and grey colored nodes show their involvement in disease resistance, defense mechanism, disease and defense mechanism, stress-related and less annotation, respectively.

### Analysis of physicochemical features of the unknown proteins

The physiochemical features of the 5 disease-resistance proteins were calculated from ExPASy’s ProtParam tool. **Table 1** shows the physicochemical features, such as atomic composition, molecular weight, theoretical isoelectric point (pI), instability index, and extinction coefficient, for all 5 proteins. The BB.2 protein has the highest number (1,212) of amino acids, whereas Protein RB.3 has the least number (487) of amino acids. Further, BB.2 has a maximum molecular weight of 134.85 kilodaltons (kDa), while Protein BB.3 has a minimum molecular weight of 50.98 kDa. The Protein RB.3 was found to have a maximum pI of 8.05, indicating that it is a positively charged protein, followed by RB.1 (pI=7.38), RB.2 (pI=7.29), BB.1 (pI=6.51), and BB.2 (pI=6.42). Protein BB.1 has the lowest instability index of 39.24, showing that it is a stable protein. Moreover, it has a high aliphatic index depicting high thermostability over a wide range of temperatures. All 5 proteins showed low GRAVY values ranging from -0.65 to 0.04. This indicates that these proteins are globular (hydrophilic) rather than membranous (hydrophobic). The extinction coefficients for the proteins range from 141,900 to 168,180 with respect to cysteine (C), tryptophan (W), and tyrosine (Y), showing the presence of high concentrations of C, W, and Y in all the proteins.

**Table 1.** Physiochemical features of five disease-resistance proteins from ExPASy’s ProtParam tool.

| Characteristics | Protein |  |  |  |  |
| --- | --- | --- | --- | --- | --- |
|  | BB.1 | BB.2 | RB.1 | RB.2 | RB.3 |
| No. of amino acids | 998 | 1212 | 674 | 1046 | 487 |
| Molecular weight (in kDa) | 108.30 | 134.85 | 73.26 | 116.54 | 50.98 |
| pI | 6.51 | 6.42 | 7.38 | 7.29 | 8.05 |
| Instability index | 39.24 | 53.57 | 52.28 | 51.40 | 54.14 |
| Aliphatic index | 104.81 | 68.81 | 59.38 | 69.82 | 55.01 |
| GRAVY | 0.04 | -0.61 | -0.72 | -0.50 | -0.65 |
| Extinction coefficients | 56350 | 141900 | 46215 | 168180 | 46215 |

Characteristics: determines physical and chemical parameters of proteins; kDa: kilodaltons; pI: isoelectric point.

### Structure prediction, refinement, and model quality assessment

PSIPRED tool predicted all the 5 proteins (i.e., BB.1, BB.2, RB.1, RB.2, and RB.3) were having a significant helix and coil content. Of these, BB.2 has the highest helix and coil content comprising 363 and 758 amino acid residues, respectively (**Table S2**). We performed homology modeling using SWISS-MODEL to predict the 3D structure of five disease-resistance proteins. We noticed that the template for RB.2 protein has the best QMQE score of 0.63, whereas the templates of RB.1 and RB.3 proteins have the best sequence identity of 84.75%. The GalaxyRefine2 server and ModRefiner were used to refine the models for all five proteins. Among the 5 proteins, BB.2 has the best model refined by the GalaxyRefine2 server. Moreover, The presence of disulphide bonds is important in stabilizing protein structures and understanding structural-functional relationships. CYSPRED showed both bonding and non-bonding state of cysteine residues. In addition, CYSPRED predicted the highest reliability score for all 5 proteins showing the significance of the prediction of the disulphide bonds.

### Stereochemical features analysis

**Table S3** shows the results from the analysis performed using the Ramachandran Plot server, PROCHECK, Verify3D, and ERRAT tools. The Ramachandran plot shows the backbone dihedral angles of proteins. The Ramachandran plot analysis shows that BB.2, RB.1, and RB.3 had the best distribution of φ and Ψ angles. In addition, for the same 3 proteins, more than 90% of amino acids were in the favored regions, and none of the amino acids were in the disallowed regions. The PROCHECK tool checks the stereochemical quality of protein structure by analyzing the residue-by-residue geometry and overall structure geometry. Likewise, Ramachandran Plot server, the PROCHECK results also showed that amino acids of BB.2, RB.1, and RB.3 proteins were in the allowed region of the Ramachandran plot. The Verify3D tool determines the compatibility of an atomic model (3D) with its own amino acid sequence (1D). According to Verify3D, the compatibility percentage was found to be 100% for BB.2 (100%), followed by BB.1 (96.58%) and RB.2 (91.11%). The ERRAT tool analyses the statistics of non-bonded interactions between different atom types. It shows that the best score out of 100 was for RB.1, followed by BB.2 (95.58) and BB.1 (94.11). The 3D models of the 5 proteins were visualized using Pymol and shown in **Figure 2**.

**Fig. 2.**
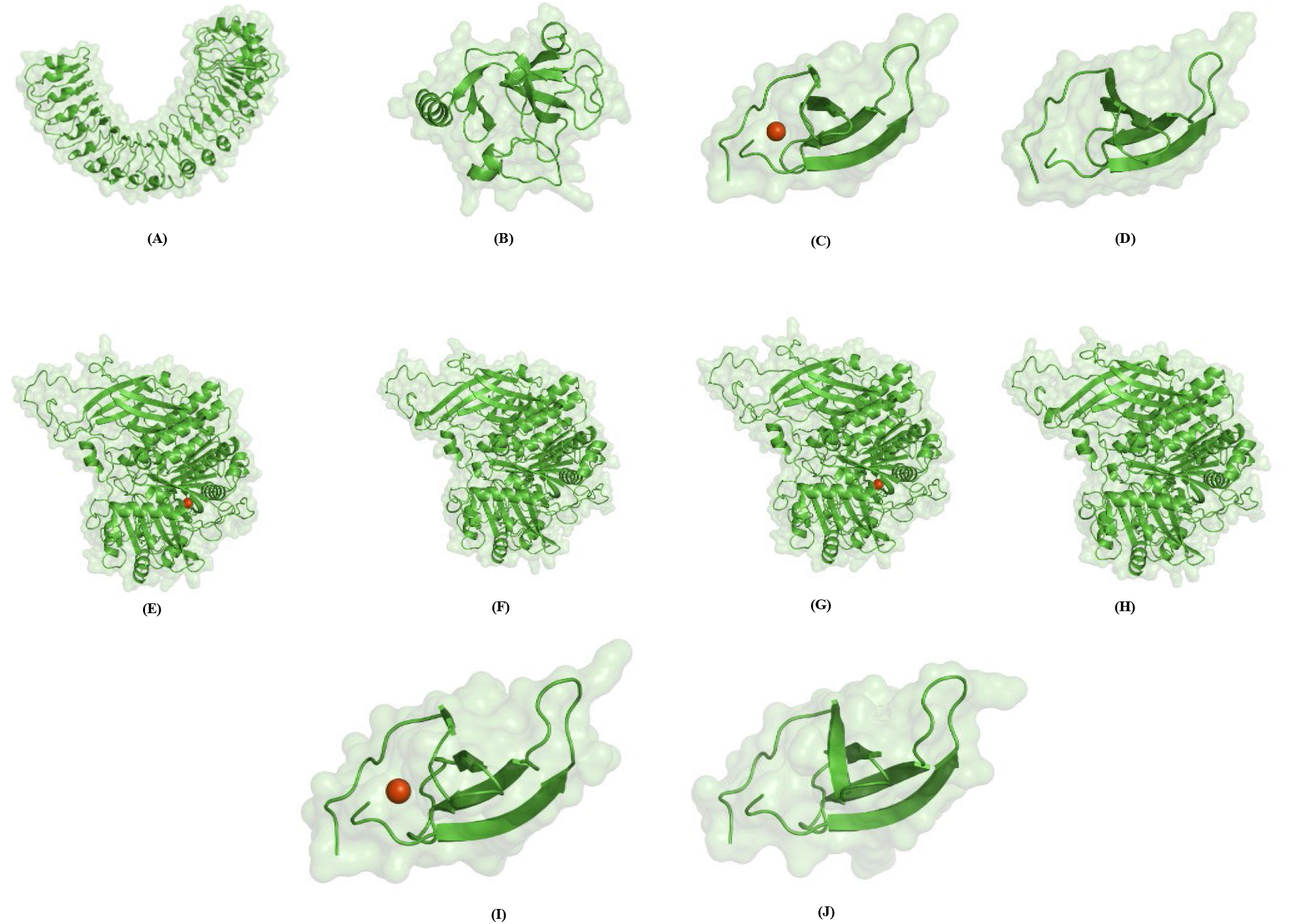
Protein models predicted by SWISS-MODEL, refined models by Galaxy Refine2 server, and ModRefiner for the proteins against bacterial blight and rice blast diseases and visualization in Pymol. **Panels (A-D)** shows predicted protein structures of Proteins BB.1 and BB.2 by SWISS-MODEL and their refined models by Galaxy Refine2 server, respectively. **Panels (E-J)** shows protein structures of Proteins RB.1, RB.2, and RB.3 by SWISS-MODEL and their refined models by Galaxy Refine2 server and ModRefiner, respectively.

### Subcellular localization and disulfide bond analysis

The subcellular localization of 5 proteins was carried out using CELLO2GO to understand their functions and role in regulating the biological processes at the cellular level. The CELLO2GO result revealed that BB.1 was the only protein found in the plasma membrane, with a score of 2.12 (42.3%). The remaining 4 proteins (i.e., BB.2, RB.1, RB.2, and RB.3) with scores of 4.42 (88.4%), 4.33 (86.6%), 2.62 (52.4%) and 3.76 (75.2%), respectively, were present in the nucleus. According to the HMMTOP tool, we found that Proteins BB.1 and BB.2 consist of two and one transmembrane helices. In addition, Protein RB.2 was predicted to contain one transmembrane helix. Moreover, we used a neural network-based predictor, CYSPRED, to predict the cysteine residues (**Table S4**). The presence of disulfide bonds is important in stabilizing protein structures and understanding structural-functional relationships. CYSPRED showed both bonding and non-bonding state of cysteine residues. In addition, CYSPRED predicted the highest reliability score for all 5 proteins showing the significance of the prediction of the disulfide bonds.

### Multiple sequence alignment and phylogenetic analysis

InterProScan and CDD tools were used for studying the domains, motifs, families, and superfamilies of the 5 unknown proteins. Three proteins, BB.1, BB.2, RB.1, RB.2 and RB.3, were found to contain the domains such as LRR, Kinase, Transmembrane (TM), Su(var)3-9, Enhancer-of-zeste and Trithorax (SET), WRKY and Phospholipase D known to be involved in disease resistance. Additionally, DRAGO3 also predicted 5 LRR, 4 Kinase, and 1 TM domains for the Protein BB.1. Similarly, from the LRR predictor, we found that there were in total 22 LRR domains in the Protein BB.1. The presence of these disease-resistant domains indicated that BB.1 might be involved in the disease-resistance mechanism against *Xoo* and *M. oryzae*. **Table 2** shows information on the conserved domains, motifs, families, and superfamilies of the 5 unknown proteins.

**Table 2.**
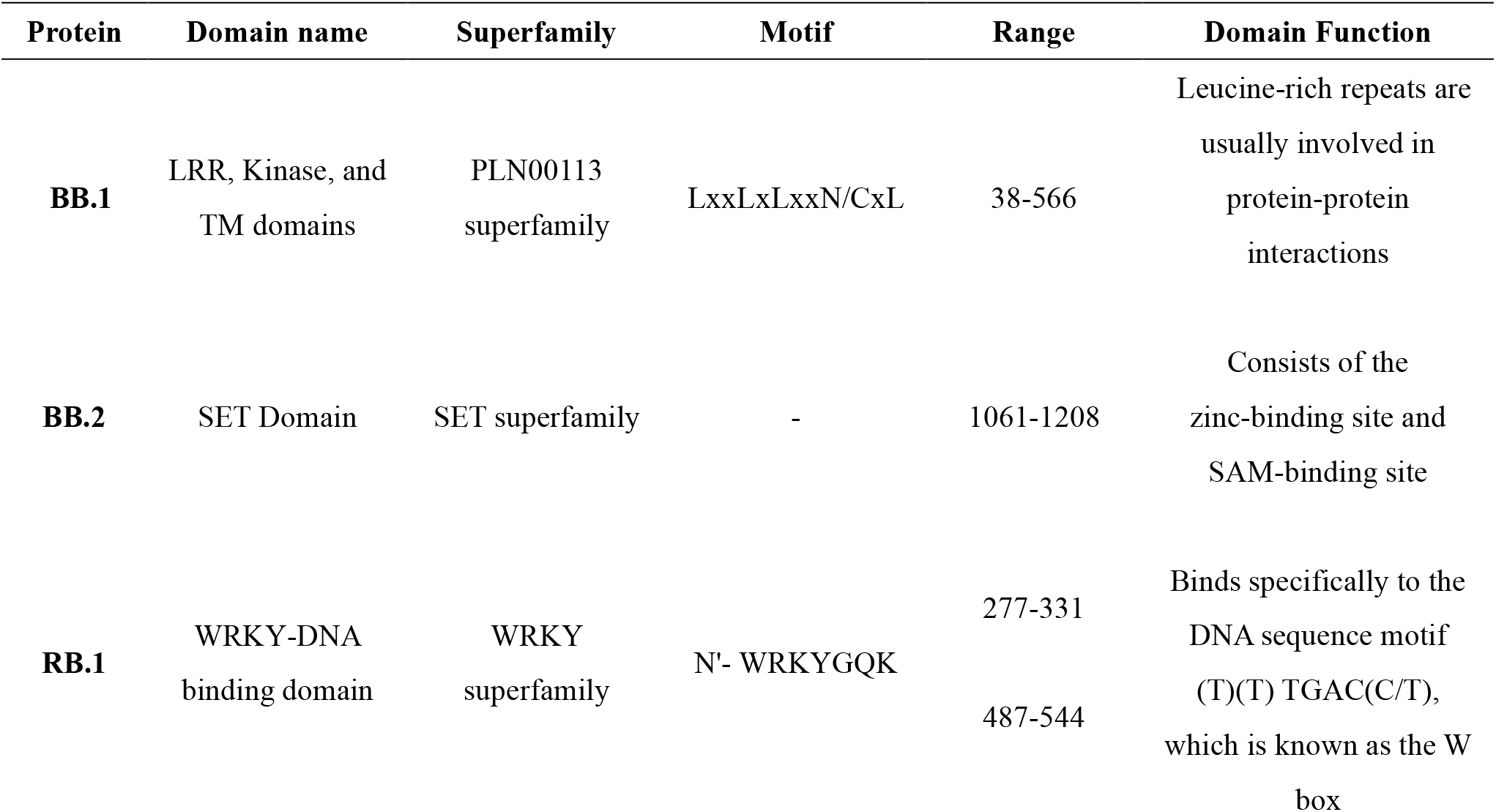
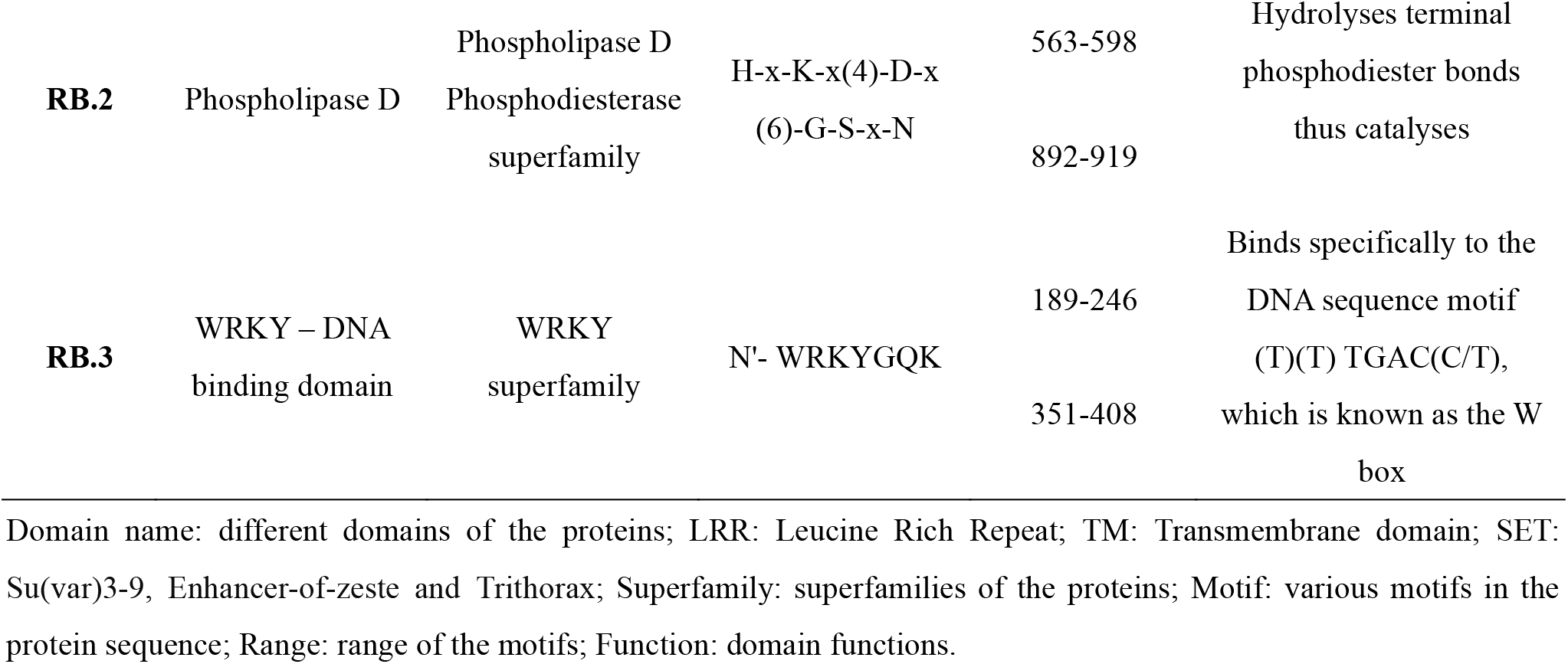
Shows the conserved domains, motifs, families, and superfamilies information of the unknown disease-resistance proteins using CDD, InterProScan, DRAGO3, and LRR Predictor.

The ScanProsite was used to identify specific motif consensus sequences of conserved domains from the iterative BLASTP search for the 5 unknown proteins. Some hundreds of proteins with identical domains and similar motif consensus were retrieved for each query motif. From these BLASTP hits, some 12 to 15 sequences were selected from related organisms either from the *Oryza* genus or *Arabidopsis thaliana* (*A. thaliana*). The above-selected sequences were found to have high similarity and annotation scores in the range of 4 to 5, as reported in the UniProtKB.

On average, we observed 70% similarity between our query proteins (BB.1, BB.2, RB.1, RB.1, and RB.3) and their respective similar proteins. For two proteins, BB.1 and RB.1, the similarity was around 80% with their similar proteins. For each of the five proteins, we performed MSA with their respective similar proteins as described above.

The phylogenetic analysis showed that the protein BB.1 was placed in the monophyletic clade with receptor kinase-like protein XA21 which is annotated with a score of 5 as per the UniProtKB (**Figure 3**). The clade bootstrapping confidence is 100, and the subsequent alignment was found to be 84% similar. Similarly, the BB.2 protein was found to be close to the Histone-lysine N-methyltransferase (OsTRX1) protein. We noticed high bootstrap scores of 100 and 98, respectively, for the protein RB.1 and RB.3, which implies that they shared a common ancestry with the conserved WRKY domain with a zinc-finger motif. This domain is commonly found in protein superfamilies involved in various plant physiological regulations, including disease resistance. Finally, the protein RB.2 shared similar phospholipase D (PLD) phosphodiesterase domain activity with its orthologs in *A. thaliana*. The active site profile in Protein RB.2 was confirmed by the presence of histidine (H), lysine (K), and aspartic acid (D) residues, which refer to the popular HKD motif consensus.

**Fig. 3.**
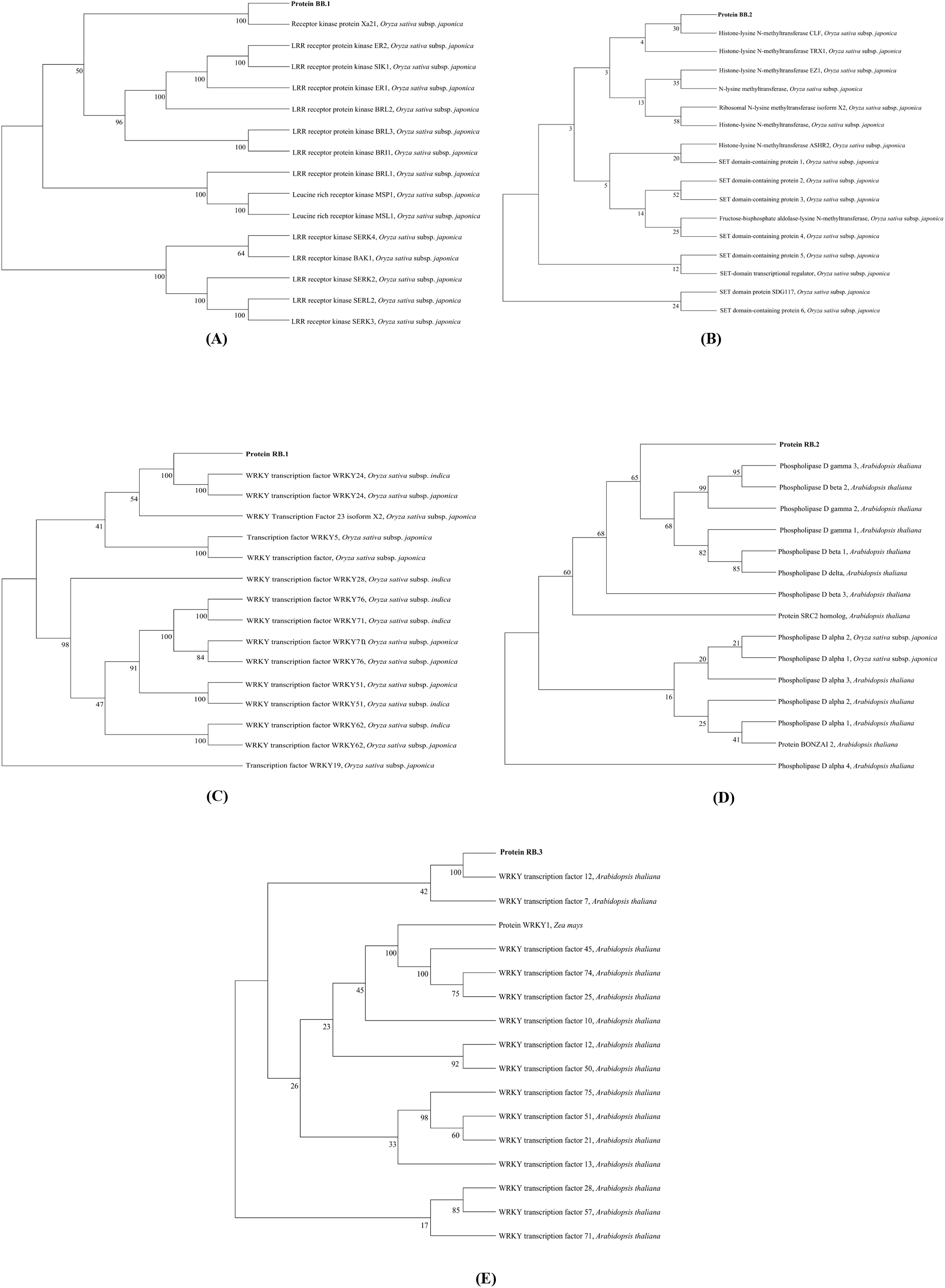
Phylogenetic trees created through Multiple Sequence Alignment (MSA) by phylogram package for the proteins against bacterial blight and rice blast diseases. (A) and (B) shows phylogenetic trees for Protein BB.1 and BB.2, respectively. (C), (D) and (E) shows phylogenetic trees for Protein RB.1, RB.2, and RB.3, respectively.

### Functional enrichment analysis

**Table 3** shows the functional enrichment analysis results using the PANNZER tool. This tool predicted that Protein BB.1, RB.1, RB.2, and RB.3 have biological functions such as protein phosphorylation, defense response, response to the bacterium, positive regulation of defense response to the bacterium, phospholipase D activity and presence of WRKY domain. This might indicate that BB.1, RB.1, RB.2, and RB.3 are involved in the disease resistance mechanism. Biological pathways analysis was performed using the Assign KO tool to find their role in the disease resistance mechanism. This tool predicted that all three proteins against RB were involved in plant-pathogen interaction, Ras, cAMP, and MAPK signaling pathways (**Table 4, Table S5**). This indicates that these proteins might be involved in the pathways responsible for providing resistance against diseases.

**Table 3.**
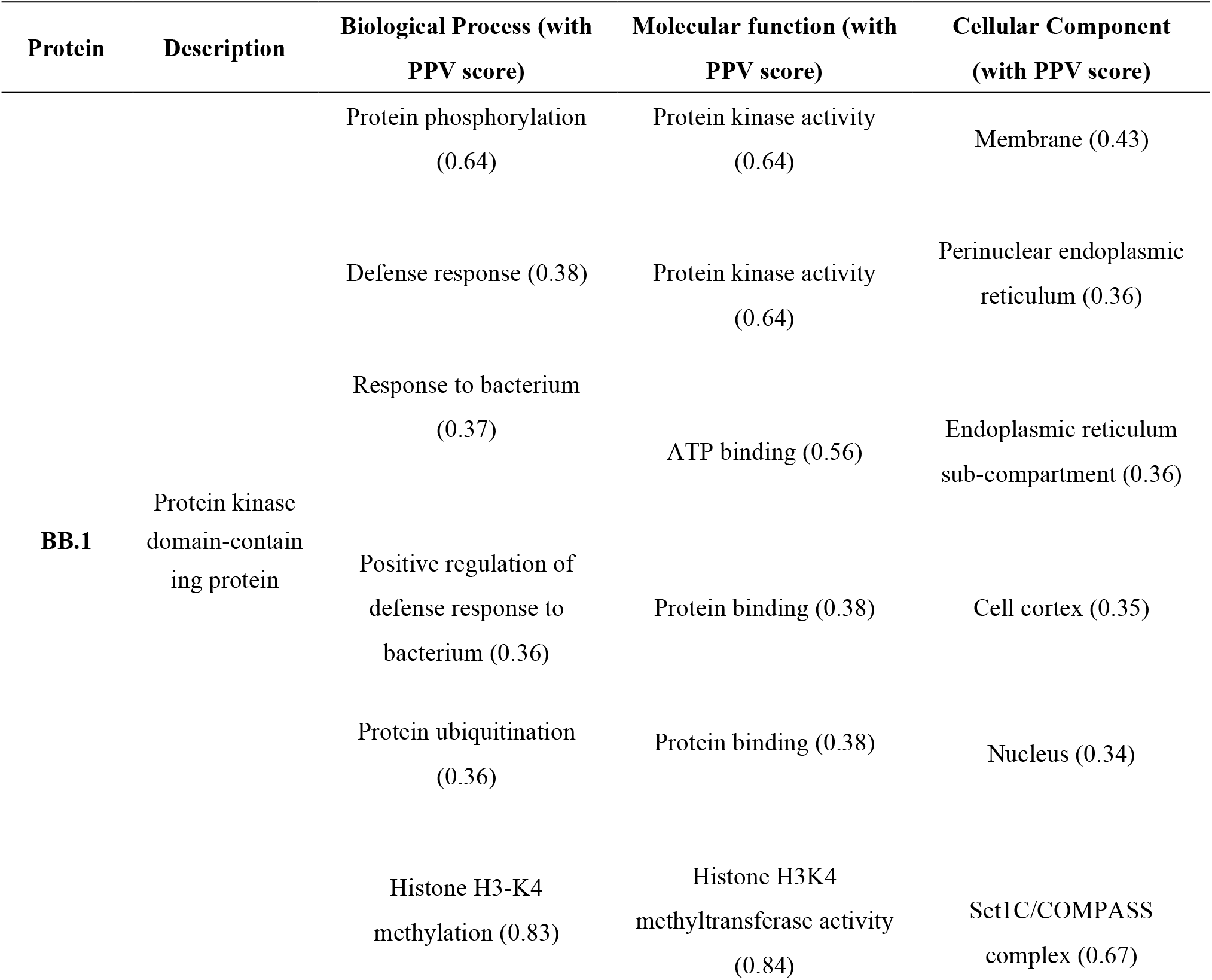
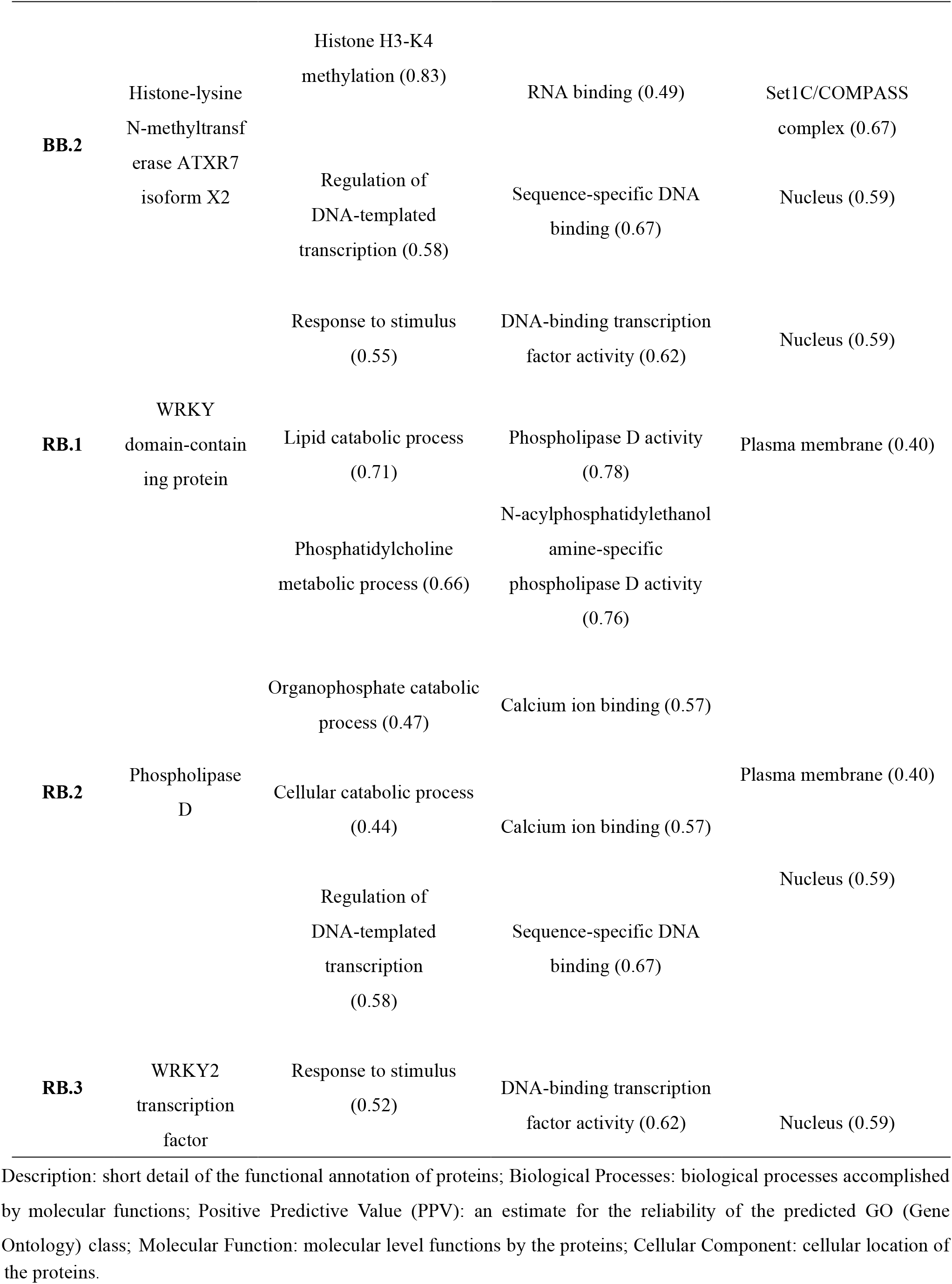
Functional annotation of the disease-resistance proteins using the PANNZER tool.

**Table 4.** Shows pathway analysis results for proteins using the Assign KO tool available in the KEGG Database.

| Protein | Pathway involved | Definition | Functional category |
| --- | --- | --- | --- |
| RB.1 | Environmental adaptation<br>(plant-pathogen interaction) | WRKY2; WRKY<br>transcription factor 2 | Organismal systems |
| RB.2 | Lipid metabolism, Ras<br>signaling pathway, and cAMP<br>signaling pathway | PLD1_2; phospholipase<br>D1/2 | Metabolism, Environmental<br>Information Processing |
| RB.3 | MAPK signaling pathway | WRKY33; WRKY<br>transcription factor 33 | Organismal systems,<br>Environmental Information<br>Processing |

Pathway involved: pathways in which proteins are involved; Definition: definition of the pathways; Functional category: functional categories of the pathways.

### Predicting functional interactors of disease-resistance proteins

The protein-protein interactions are of central importance for every process in a living cell. The Expasy STRING database indicated the interactions between transcription initiation factor IIA subunit 2 (Q0DLD3, TFIIAy) and WRKY transcription factor WRKY24 (Q6IEQ7, WRKY24) with the Protein BB.1. In addition, we identified the interaction of Protein RB.3 with mitogen-activated protein kinase 1 (Q84UI5, MPK1).

## DISCUSSION

Food security is a global concern, especially for staple crops such as rice which prompts a greater focus on developing and improving crop protection methods. The regular increase in the application of chemical fertilizers and pesticides in rice has made the pathogens more resistant, and hence, there is a global effort for pest biocontrol including the development of disease-resistant varieties for future global food security. This work aimed to identify unknown resistance proteins against major bacterial and fungal diseases in *Oryza sativa* sp. *japonica* and their *in silico* characterization using state-of-the-art bioinformatics tools. We focused on the disease-resistance proteins for BB and RB diseases following a genome-wide study. We identified five unknown disease-resistance proteins, two for BB and three for RB, and characterized them according to their functions, domains, protein-protein interactions, and involvement in biological pathways.

Gene co-expression and protein-protein interaction network analyses are powerful *in silico* tools to identify unknown candidate genes (Klasberg et al., 2016). Firstly, we collected data for known and disease-resistance genes from Ensembl plants and SNP-Seek databases. Next, we performed a network-based analysis to identify unknown disease-resistance proteins in rice. Thus, for BB disease, network analysis using Cytoscape revealed 17 disease-resistance genes, five defense-related genes, and five genes involved in disease resistance and defense mechanisms. Similarly, for RB disease, we identified 11 genes for disease resistance, eight genes as defense-related, and seven genes involved in both disease resistance and defense mechanisms. After this, we performed various in-silico analyses of these five identified proteins. MSA and phylogeny analyses were performed, which showed the evolutionary relationship of the predicted proteins with reported disease-resistance proteins. In addition, secondary and tertiary structures were predicted and validated along with the stereochemical and physicochemical properties. The subcellular localization revealed their presence in the nucleus and plasma membrane, whereas the topology study indicated the presence of transmembrane helices in them. Furthermore, the biological sequence analysis, including disulfide bonds, conserved domains, motifs, folds, families, and superfamilies, was performed, revealing structural features of plant disease resistance proteins.

We found that BB.1, BB.2, RB.1, RB.2, and RB.3 carry LRR, Kinase, TM, SET, WRKY, and Phospholipase D domains known to be involved in disease resistance against *Xoo* and *M. oryzae*. Our results pointed at a high similarity of Protein BB.1 with Rice Receptor Kinase XA21 (RRK XA21), which indicates a similar immune response action against BB disease. Park & Ronald (2012) have found that the RRK XA21 is responsible for broad-spectrum innate immunity against the bacterial pathogen *Xoo*. During bacterial infection, RRK XA21 recognizes the *Xoo* Ax21 protein and binds to its conserved sulfated peptide region to finally release a kinase domain upon cleavage, and this kinase domain ultimately translocates into the rice protoplast nucleus. Following the nuclear localization, the XA21 kinase domain binds with transcriptional regulation factors to trigger the immune response.

Furthermore, Choi et al., 2014 experimentally validated the function of OsTRX1, which contains a SET domain similar to our Protein BB.2. The SET domain is essential for the methylation of a lysine residue of the N-terminus of Histone H3 (Rea et al., 2000). In rice, this methylation is crucial for the OsTRX1-mediated negative regulation of transcriptional activation, which is part of a broader defense mechanism against BB disease. Protein BB.2 and OsTRX1 share a matched lysine residue in their respective SET domains, followed by a high cystine signature in the post-SET region towards the C-termini. Thus, the prediction of methyltransferase activity in Protein BB.2 is supported by its similarity with high cystine and lysine-rich consensus in OsTRX1. Moreover, proteins RB.1 and 3 RB.3 shared a common ancestry with the conserved WRKY domain with a zinc-finger motif, which is commonly found in protein superfamilies involved in various plant physiological regulations, including disease resistance. The high confidence score of proteins in subsequent phylogenetic analysis indicates a strong probability of their functioning as defense-related transcriptional regulators. In addition, RB.2 shared a similar phospholipase D (PLD) phosphodiesterase domain with its orthologs in *A. thaliana*. The active site profile in RB.2 was confirmed by the presence of histidine (H), lysine (K), and aspartic acid (D) residues, which constitute the well-known HKD motif consensus. Our phylogenetic study illustrated the common ancestry of HKD motifs presented in various copies of phospholipase phosphodiesterase proteins of *A. thaliana*. These paralogs are known candidates in the resistance mechanism of *A. thaliana* against RB disease (Bargmann & Munnik, 2006). Therefore, from a strong homology perspective, RB.2 is predicted to possess similar hydrolyzing activity of glycerol phospholipids during the resistance against the RB disease in rice.

Functional enrichment analysis of the predicted proteins showed their involvement in the disease resistance mechanism. BB.1, RB.1, RB.2, and RB.3 were found to have roles in biological functions such as protein phosphorylation, and phospholipase D activity. In addition, BB.2 was predicted to be involved in Histone H3-K4 methylation, RNA binding, and DNA-binding transcription factor activity. The pathway analysis of BB.1 and BB.2 has also indicated their roles in transcription under control conditions as well as bacterial infection. Moreover, RB.1 was involved in secondary metabolite synthesis and lipid metabolism, whereas RB.2 and RB.3 were involved in Ras, phospholipase D, cAMP, and MAPK signaling pathways, indicating their possible functional mechanisms in the resistance mechanism against *Xoo* and *M. oryzae*. Furthermore, the functional interactors of our identified proteins were also found to be involved in disease resistance. For example, the interactors of BB.1 and RB.3, viz. TFIIAy and WRKY24, respectively, are known players of bacterial disease resistance in rice and are highly expressed in disease-resistant rice varieties (Kumar et al., 2022; Zhang et al., 2017). This strongly suggested that our identified proteins are likely to be involved in rice disease resistance against RB and BB.

In summary, the current study predicted five unknown disease-resistance proteins in rice against *Xoo* and *M. oryzae*. Future experimental validation of the predicted roles of these proteins will provide new candidates for disease resistance in rice. This will help rice breeders and biotechnologists for developing resistant rice varieties against BB and RB diseases. This information will also be used to develop molecular markers that can be used to screen varieties for resistance to both BB and RB diseases, as well as for marker-assisted selection.

## Author Contributions

VD collected study data, performed primary analysis, and designed the working strategy; SB carried out basic data analysis and contributed to manuscript writing; RSS performed functional enrichment; PY conceptualized and supervised the study; AS edited the manuscript and provided inputs; all authors contributed to writing the manuscript.

## ACKNOWLEDGEMENTS AND FUNDING

PY acknowledges the support from seed grant (project number I/SEED/PY/20200037) funded by the Indian Institute of Technology, Jodhpur, India; VD is thankful for the support from the Ministry of Human Resource and Development (MHRD), India fellowship and RSS (file number:09/1125(0019)/2021-EMR-I) is supported by the CSIR-NET fellowship.

## CONFLICT OF INTEREST

Authors declare that there are no competing financial interests.

## SUPPLEMENTAL DATA

The supplemental data is available in the online version of this article.

## Notes

### Competing Interest Statement

The authors have declared no competing interest.

