## SUPPLEMENTARY DATA for "*In silico* characterization of five novel disease-resistance proteins in *Oryza sativa sp. japonica* against bacterial leaf blight and rice blast diseases"

**Correspondence**:

Pankaj Yadav

Assistant Professor

**Supplementary Tables**

**Table S1.** Overview of different databases and tools used in the study.

| Categories | Description |
|---|---|
| **Databases** | |
| SNP-Seek database | Database providing information of genotype, phenotype, and variety for rice derived from 3,000 Rice Genomes Project |
| Expasy STRING (Search Tool for the Retrieval of Interacting Genes/Proteins) | Knowledgebase and software tool for protein-protein interactions |
| UniProt Knowledgebase (UniProtKB) | Richly curated protein database having protein sequence and functional information |
| ModelArchive | Archive for structural models which are not based on experimental data and are computationally derived |
| Conserved Domain Database (CDD) | Resource for the annotation of protein sequences based on the location of conserved domain footprints |
| DRAGO | Web-based tool in Plant Resistance Genes Database (PRGdb) database for providing a comprehensive overview of resistance genes (R-genes) in plants |
| STRING database | Database of known and predicted protein-protein interactions |
| **Tools** | |
| Biomart tool | Open-source data management webtool for data refinement and retrieval available in Ensembl plants database |
| Cytoscape | Open-source software platform for visualising of biomolecular interaction networks |
| PSI-BLAST Based Secondary Structure Prediction (PSIPRED) | Protein Analysis Workbench/ tool for prediction of secondary structure |
| SWISS-MODEL | Web-based protein structure homology-modelling server |
| GalaxyRefine2 server | Web server for protein structure prediction, refinement, and related methods |
| ModRefiner | Algorithm for atomic-level, high-resolution protein structure refinement |
| Pymol | Open-source molecular visualization system for biomolecules especially proteins |

| | |
|---|---|
| Ramachandran plot server | Web-based server for constructing Ramachandran plot |
| PROCHECK | Web-based tool for checking the stereochemical quality of a protein structure |
| Verify3D | Web-based tool for determining the compatibility of an atomic model (3D) with its own amino acid sequence (1D) |
| ERRAT | Web-based tool for evaluating protein models |
| ExPASy ProtParam | Web-based tool that computes various physical and chemical parameters for a given protein |
| CYSPRED | Neural-Network-based predictor for predicting bonding state of cysteines in proteins |
| ModelArchive | Archive for structural models which are not based on experimental data and are computationally derived |
| CELLO2GO | Web server for protein subcellular localization prediction with functional gene ontology annotation |
| HMMTOP | Automatic server for predicting transmembrane helices and topology of proteins |
| InterProScan | Web-based tool for functional analysis of proteins by classifying them into families and predicting domains |
| ScanProsite | Web-based tool for detecting PROSITE signature matches in protein sequences |
| Motif Scan | Finds all known motifs occurring in a protein sequence |
| *ggmsa* package (R software) | Tool for visualization and annotation of multiple sequence alignment of protein/DNA/RNA |
| *phylogram* package (R software) | Tool for developing phylogenetic trees |
| PANNZER (Protein Annotation with Z-score) | Webserver for functional annotation of proteins of unknown function |
| Assign KO tool (KEGG Database) | Interface to the BlastKOALA server for pathway analysis |
| Leucine Rich Repeats (LRR) predictor | Webserver for predicting leucine-rich repeats |

Categories: tools and databases used in the study; Description: short detail of the tool and databases used.

**Table S2.** The secondary structure profile of the five unknown disease-resistant proteins.

| Structure | Number of amino acid residues |
|---|---|

| | BB.1 | BB.2 | RB.1 | RB.2 | RB.3 |
|---|---|---|---|---|---|
| **Helix** | 280 | **363** | 94 | 191 | 61 |
| **Coil** | 603 | **758** | 554 | 680 | 400 |

Structure: number of helices and coils present in the protein; BB.1 and BB.2: disease-resistant protein against bacterial blight disease; RB.1, RB.2 and RB.3: disease-resistant protein against rice blast disease.

**Table S3.** The stereochemical analysis of the five disease-resistant proteins using different tools.

| Proteins | Ramachandran Plot | PROCHECK | Verify 3D | ERRAT |
|---|---|---|---|---|
| **BB.1** | 0.7% | 0.8% | 96.58% | 94.11 |
| **BB.2** | 0% | 0% | 100% | 95.58 |
| **RB.1** | 0% | 0% | 80.70% | 100 |
| **RB.2** | 0.1% | 0.3% | 91.11% | 79.63 |
| **RB.3** | 0% | 0% | 83.05% | 84.09 |

Protein: five disease-resistant proteins; PROCHECK: checks the stereochemical quality of protein structure by analyzing residue-by-residue geometry and overall structure geometry; Ramachandran Plot: refers to the percentage of amino acids in the disallowed region; Verify 3D: compatibility percentage of an atomic model (3D) with its own amino acid sequence (1D); ERRAT: depict the score (out of 100) of the non-bonded interactions between different atom types.

**Table S4.** Shows CYSPRED predicted cysteine residues playing important in disulphide bonding.

| Proteins | Number of cysteine residues | Predictions | Reliability score |
|---|---|---|---|
| **BB.1** | 21 | Bonding state (3), non-bonding state (18) | 9 |
| **BB.2** | 30 | Non-bonding state | 9 |
| **RB.1** | 8 | Non-bonding state | 9 |
| **RB.2** | 14 | Non-bonding state | 9 |
| **RB.3** | 8 | Non-bonding state | 9 |

Number of cysteine residues: total number of cysteine residues present in the proteins; Predictions: bonding and non-bonding state of the cysteine residues; Reliability score: score for the reliability of the predictions.

**Table S5.** Information on pathway analysis of the disease-resistant proteins.

| Proteins | Pathways involved | Definition | Functional category |
|---|---|---|---|
| **BB.1** | Transcription (Basal transcription factors) | TFIIB, GTF2B, SUA7, tfb; transcription initiation factor TFIIB | Genetic Information Processing |
| **BB.2** | Transcription (Basal transcription factors) | TAF5; transcription initiation factor TFIID subunit 5 | Genetic Information Processing |
| **RB.1** | Environmental adaptation (plant-pathogen interaction) | WRKY2; WRKY transcription factor 2 | Organismal systems |

| | | | |
|---|---|---|---|
| **RB.2** | Metabolism (secondary metabolites biosynthesis, Glycerophospholipid and ether lipid metabolism), Environmental Information Processing (Ras, phospholipase D, sphingolipid, cAMP signaling pathways) | PLD1_2; phospholipase D1/2 | Lipid metabolism |
| **RB.3** | Environmental Information Processing (MAPK signaling pathway), Environmental adaptation (plant-pathogen interaction) | WRKY33; WRKY transcription factor 33 | Environmental Information Processing |

Proteins: five disease-resistant proteins; Pathways involved: details of the identified pathways; Definition: short details of the disease-resistant proteins; Functional category: description of the biologically identified functional groups in the proteins.
